# Mutational sequencing for accurate count and long-range assembly

**DOI:** 10.1101/149740

**Authors:** Vijay Kumar, Julie Rosenbaum, Zihua Wang, Talitha Forcier, Michael Ronemus, Michael Wigler, Dan Levy

## Abstract

We introduce a new protocol, mutational sequencing or muSeq, which randomly deaminates unmethylated cytosines at a fixed and tunable rate. The muSeq protocol marks each initial template molecule with a unique mutation signature that is present in every copy of the template, and in every fragmented copy of a copy. In the sequenced read data, this signature is observed as a unique pattern of C-to-T or G-to-A nucleotide conversions. Clustering reads with the same conversion pattern enables accurate count and long-range assembly of initial template molecules from short-read sequence data. We explore count and low-error sequencing by profiling a 135,000 fragment PstI representation, demonstrating that muSeq improves copy number inference and significantly reduces sporadic sequencer error. We explore long-range assembly in the context of cDNA, generating contiguous transcript clusters greater than 3,000 bp in length. The muSeq assemblies reveal transcriptional diversity not observable from short-read data alone.

## INTRODUCTION

Long-read sequencing platforms such as PacBio and Oxford Nanopore are costly and error-prone, but provide the long-range information required for high quality assemblies (1). Short-read sequencers are relatively inexpensive and have excellent precision; however, the reads lengths are sufficient only for simpler assemblies. We tested a theoretical idea, random template mutagenesis, which bridges this gap: it converts short-read sequencers into virtual long-read sequencers. Greater precision and accurate counting are added benefits of the method.

The specific problems with short-read sequencers are readily enumerated. Whenever the distinguishing variants in the template molecules are more than one read length apart, multiple distinct assemblies are equally consistent with the read data. This prevents resolving haplotypes, observing transcript isoforms, and assembling complex repetitive regions. Although sequence fidelity is good, low-frequency variants are not distinguishable from PCR and sequencing error. Finally, distortion during PCR amplification makes for an unreliable estimate of count in RNA expression and DNA copy number.

Recently, we proposed a theoretical solution to these problems (2). By marking each initial template molecule with a random mutation pattern, all subsequent copies of the original molecule will carry the same pattern. This enables both accurate counting and low-error sequencing. Further, overlapping copies from the same initial template can be joined computationally if they have compatible patterns that far exceed chance agreement. With sufficient coverage, this property would enable the long-range assembly of each mutated template molecule. Such information is also useful for problems of haplotype phasing, measuring repeats, and detecting rare variants with confidence.

Here we introduce an implementation of this idea, which we call mutational sequencing or muSeq. We use partial sodium bisulfite conversion to mark template double-stranded DNA molecules or first-strand cDNAs. The bisulfite reaction deaminates unmethylated cytosines, and it is typically used for studying cytosine methylation patterns in the genome (3). In that application, the deamination reaction is run to completion, converting nearly every unmethylated cytosine to uracil. For randomly marking templates, however, we require partial conversion. By adjusting the time and temperature of a step in the bisulfite reaction, we can reliably control the rate of conversion. Reflecting the binary nature of the conversion, we refer to cytosines in this context as “bits.”

To test the operating characteristics of the muSeq protocol, we conducted two series of experiments. In the first we studied the application of muSeq to a genomic representation. Our experiments show that the rate of deamination is independent of position and uncorrelated within the template. We observe that fragment counts are linear with copy number and that allele ratios follow the expected binomial distribution. We determine that the method does not contribute any measurable sequence error. In the second series of experiments, we applied the muSeq protocol to cDNA derived from reverse transcribed poly(A)^+^ cellular RNA. Applying a simple algorithm, we clustered sequence reads into a longer consensus template. The resulting consensus templates compare favorably to reference transcript assemblies. Analyses of the data demonstrate the ability to reconstruct splicing patterns at the level of individual transcripts.

### MATERIALS AND METHODS

#### Representations

Genomic DNAs were extracted from whole blood, cleaved with PstI, end-repaired and ligated to custom Illumina sequencing primers. The primers are rendered conversion-resistant by substituting 5-methylcytosine (5mC) for cytosine during oligo synthesis. The complete conversion protocol uses the MethylEasy Xceed Rapid DNA Bisulphite Modification Kit Mix (Human Genetic Signatures/Clontech) according to standard instructions. The partial conversion protocol uses the same kit and instructions, but we reduce the temperature and time during incubation with the combined Reagents 1 and 2 (step 5 in the instructions). We started with 75 ng of input DNA for each reaction and carried out both complete (45 min, 80°C) and partial conversion at 3, 6 and 9 minutes at 73°C. After conversion, we sampled 4% from each converted sample and PCR-amplified using Illumina P5 and P7 sequencing adapters (for the complete conversion library, we sampled 40%). The resulting libraries were sequenced on an Illumina MiSeq (~17 million paired-end reads per sample). We also sampled 2% from the 6 minute conversion, then amplified for five linear rounds with just one primer (P7); we then completed the PCR as above, sequencing the resulting libraries on two lanes of an Illumina NextSeq (~800 million paired-end reads). All sequencing on Illumina instruments was in paired-end 150-bp read mode, except where stated otherwise.

The conversion process operates on single stranded molecules and as such, we distinguish the two strands by their orientation and sequence as “reference top” (RT) or “reference bottom” (RB), adapting earlier usage (4). Because the sequencing adapters are attached asymmetrically, the initial template strand is always read 1 and its complement is read 2. Consequently, the conversions read by the sequencer should appear as C-to-T conversions in read 1 and G-to-A conversions in read 2. Adapting the approach of the Bismark mapper for bisulfite data (4), we first generate auxiliary read files that convert all C to T in read 1 and all G to A in read 2. These modified reads are then mapped by Bowtie2 (5) to two modified versions of the reference genome (hg38 assembly): one with all C converted to T (hg38_CT), and one with all G converted to A (hg38_GA). Selection of the best mapping determines the strand of origin. By referencing the original read pair, we determine the conversion pattern.

From the reference genome, there are 162,353 expected PstI fragments between 150–400 bp in length. We convert these fragments in silico, both C-to-T and G-to-A, and map them to hg38_CT and hg38_GA. Those in which both top and bottom strands map unambiguously (MAPQ ≥40) comprise 135,262 high quality representation fragments (HQRFs). We further consider only those reads that map with high quality alignments to HQRFs. These reads account for about 50% of the raw sequence.

Read pairs are binned by restriction fragment and the RT or RB of the initial template. Each bin is analyzed separately to determine the set of initial template conversion patterns. While many read pairs from the same initial template fragment bear identical conversion patterns and sequence, sequencing and PCR errors are sufficiently frequent to require methods for inferring their consensus, or common ancestral template. Consequently, we extract all bits from each read pair, establishing a bit string where 0 indicates that a position is unconverted and 1 indicates that a position is converted by sodium bisulfite. To cluster read pairs, we use transitive propagation (6), an algorithm we developed to find an optimal clustering. Given a model for base-calling error and a model for conversion rates, transitive propagation identifies a clustering solution that optimizes pairwise probabilities of belonging to the same cluster (a = b) or not (a ≠ b) under the condition that belonging to the same cluster is a transitive relation.

#### cDNA

We extracted total RNA from 3 million fibroblasts from a line derived from the same donor as the whole blood sample. We sampled 3.3% of the RNA for conversion to cDNA by reverse transcriptase (100 U; SMARTScribe reverse transcriptase; Clontech), employing custom oligo d(T) primers and template switch primers, each with a sample tag and random barcode. We made two such samples with distinct pairs of sample tags. We subjected the first strand cDNA to 6-minute partial bisulfite conversion as above. We selected 2.5% from each sample, amplified by PCR, mechanically fragmented (Covaris), end-repaired, adapted for sequencing with distinct library tags, and amplified. The two libraries were pooled and sequenced in two runs on a MiSeq (20 million reads).

Reads were mapped to the genome much as described above with a few key differences. First, primer and barcode sequences are trimmed from the reads (if present). Second, because sequencing adapters are added after the library is amplified, we cannot know if the conversions are C to T on read 1 and G to A on read 2, or the reverse. Consequently, we have four mappings to consider: two read conversions to two genome conversions. Finally, because Bowtie2 is not designed for mapping cDNA, we employed the STAR mapper (7). Reads that map to hg38_CT (RT reads) correspond to transcripts that match the opposite (minus) strand. Conversely, RB reads correspond to the plus strand. For a reference transcriptome, we used GENCODE release 21 (8). We restrict our attention to reads that have at least 20 base pairs mapped to annotated transcription positions on the correct strand.

We first split reads by whether they map to RT or RB. We then partition those reads into connected components: two reads are in the same component if they overlap at one or more positions, or if they are from the same read pair. Within a partition, our initial clusters are comprised of read pairs. Then, restricting to bit positions in annotated genes on the proper strand, we record matches and mismatches of bits between overlapping clusters. From these values, we apply the same noise and conversion model as in the representation to measure the pairwise probability that the clusters derive from the same initial template (blue edge) or different initial templates (red edge). Starting with the most confident pairs (for either red or blue), we add edges that do not conflict with current information and do not violate the transitivity of blue edges and stop when pairs cease to be confident (probability >1 in 10^-4^). Clusters joined by blue edges are then merged into a single new cluster.

Large partitions with many pairwise comparisons can push the limits of memory. In those cases, we subdivide the partition, consolidating no more than 1000 clusters at a time and iterating until the number of clusters is unchanged. Finally, to minimize false joins, each cluster is then tested in isolation and subjected to clustering as though within its own partition. For each cluster of reads, we record information including the most common 5′ and 3′ tags, the counts of those tags, the total number of tags for each end of the molecule, consistency of tag orientations with a transcript, positions covered, and the number of reads with each library tag.

## RESULTS

### Conversion and clustering for genomic representations

We first chose to test muSeq on genomic DNA. Obtaining high depth of coverage on the entire genome over many templates would make the study expensive to perform. We therefore chose to reduce sequence complexity by making genomic representations, in which short restriction fragments are selected and amplified to create a reproducible subset of the genome. The adapters are ‘bisulfite-resistant,’ i.e. with all cytosines methylated. The representation is comprised of many distinct sets of identical templates that are indistinguishable without bisulfite conversion. This allows us to measure conversion rates at many identical loci and within many identical templates. We chose a male donor so we could examine copy number by comparing the X chromosome to the autosomes. We also selected one particular donor for which we had whole blood for extracting genomic DNA, fibroblast cell lines for preparing RNA, and the complete genomic sequence with haplotype phasing from family information and single-cell sperm sequencing.

Based on initial results with PhiX as a template, we derived conditions for partial bisulfite conversion, and tested these on PstI representations at incubation times of 3, 6, and 9 minutes at 73°C. We compared this to the standard conditions for full conversion, 45 minutes at 80°C (Figure 1A). After making libraries, sequencing, and mapping, we examined the frequency of conversion by position and template. The three time points show a mean conversion rate of 19, 41 and 55% per template, respectively, demonstrating that the conversion rates over this range are roughly linear with time. We chose the six-minute incubation for the remainder of our experiments.

**Figure 1.**
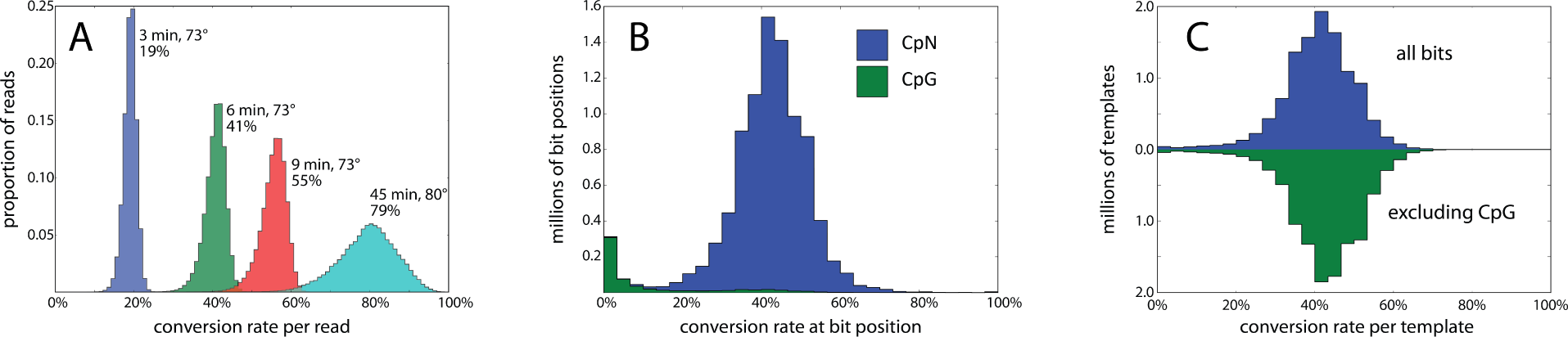
Sodium bisulfite conversion rates. **(A)** Using a kit for sodium bisulfite conversion (Materials and Methods), the standard protocol (cyan) involves a step with a 45-minute incubation at 80°C. This converts (on average) 79% of cytosines, consistent with expected rates of genomic cytosine methylation. Reducing the temperature to 73°C and the time to 3, 6 and 9 minutes results in mean conversions of 19%, 41% and 55%, respectively. **(B)** A high-depth sequencing library from the 6-minute conversion shows a mean conversion rate 42% per cytosine position with near zero conversion of cytosines in the CpG context. **(C)** The per-template cluster mutation rate is shown as a histogram. The majority of template clusters are consistent with independent conversion at a fixed rate, whether or not we exclude CpG from the count. However, there are some templates—about 0.5%—that largely escape conversion.

We made a deep-coverage library over an estimated 100 templates at an average of 30 reads per template. We first partitioned reads by mapping to PstI fragments in the expected length range. Within a partition, we clustered the reads by conversion patterns alone using transitive propagation (6). We determined that the clusters had correctly aggregated reads by template by analyzing the known heterozygous sites. >99% of clusters covering a heterozygous site had almost exclusively sequence reads with only one of the two alleles (see also section on sequence fidelity). This process is illustrated in Figure 2. Panel A shows the first 60 reads for a single, arbitrary restriction fragment in IGV (9). A single heterozygous site is indicated in the figure. Panel B shows these same reads re-ordered and grouped by conversion pattern. The heterozygous site segregates by cluster: every read in cluster 1 shows the T allele, whereas every read in cluster 2 shows the C allele. In panel C, a ‘collapsed view’ brings about 40 clusters into view, each composed of approximately 30 reads; within a cluster, all reads report the same base at the heterozygous locus.

**Figure 2.**
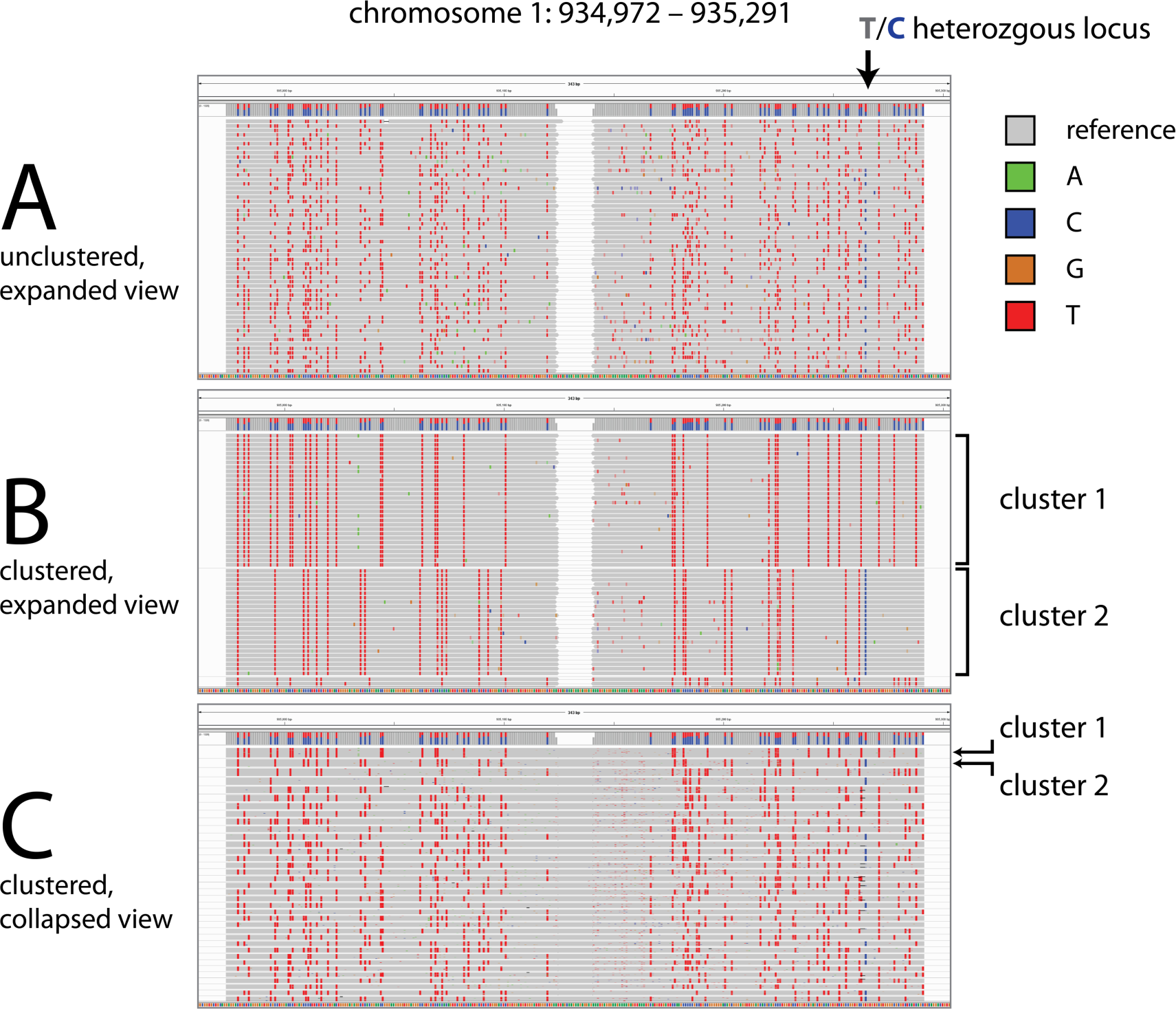
Read clustering by sodium bisulfite conversion pattern. We display IGV screenshots of muSeq reads from a 320-bp restriction fragment from chromosome 1. (**A and B)** show an expanded view of the first 60 read pairs mapped to the reference-top (RT) strand of this restriction fragment. Nucleotides matching the reference sequence are grey, with differences marked in colors reflecting the base. Notably, the red marks are homozygous C positions that converted to T. An arrow marks a heterozygous T/C position in our sample genome that is recorded as a T in the reference genome (grey) and the C allele is shown in as a blue mark. **(A)** Initially, the reads are randomly ordered. **(B)** We then cluster the reads by transitive propagation (Materials and Methods) to recover the initial template sequence. The first cluster includes 30 reads with nearly identical conversion patterns. **(C)** We show a collapsed IGV view of the first 37 clusters, each comprised of about 30 reads. Clusters 1 and 2 are indicated in both panels. The heterozygous T/C position is observed as either all T or all C in every cluster.

### Conversion follows a random uniform distribution

We used the deep coverage library to characterize randomness and independence of conversion events. We call a template ‘well-covered’ if it is supported by at least 10 reads, and we call a position ‘well-covered’ if it is covered by at least 20 well-covered templates. Of the 9 million well-covered, homozygous bit positions, 580,000 are CpG dinucleotide motifs that are predominately unconverted and account for nearly all unconverted positions (Figure 1B). For each of 11 million well-covered templates, we computed the proportion of bits converted in the template with and without CpG bits (Figure 1C, blue and green respectively). There is a marked overabundance of under-converted template, with 0.5% of templates showing fewer than 1 in 20 bits converted. Excluding these poorly converted templates, conversion rates per template are well-approximated by a fixed, independent probability of conversion for each non-CpG bit.

Moreover, conversion patterns are uncorrelated, either by position or template. To demonstrate this, we first looked at a randomly selected restriction fragment (Supplementary Figure 1A) and considered a pair of bit positions (yellow arrows). In each template of that fragment, either both bits are unconverted (0,0), both bits are converted (1,1), or only one of the two bits is converted (0,1 and 1,0). Counting these values over each of these templates produces a 2x2 contingency table (Supplementary Figure 1B, yellow box), which yields a probability that the observed counts are independent (using Fisher’s exact test). We computed this probability over all pairs of bit positions for 1000 randomly selected fragments, accounting for 2.9 million pairwise comparisons. The empirical distribution of Fisher exact p-values corresponded to the theoretical expectation assuming all pair comparisons are independent, as is seen in a Q-Q plot (Supplementary Figure 1C, circles).

Similarly, we tested whether two templates from the same fragment have independent bit patterns by looking at conversions across templates (the green arrows, boxes, and crosses in Supplementary Figure 1). In both cases, the Q-Q plots were nearly linear, suggesting that the observed distributions do not diverge for the null expectation and that deamination events are random and independent. The small deviation from expectation may reflect CpG methylation, which tends to occur in clusters. We conclude that the conversion rate is easily controlled, and the conversions themselves are independent and random.

### Counting the numbers of templates

Because our representation is drawn from a male, we can readily observe copy number difference by comparing the X chromosome with the autosomes. In a representation, read counts per fragment vary widely, reflecting varying rates of amplification due to restriction fragment length and base composition. However, if we exclude fragments containing heterozygous fragment lengths, fragments with low coverage, and fragments with very different counts for the top and bottom strands, we find that template counts accurately reflect copy number. Supplementary Figure 2A shows the distribution of template counts over the autosomes and X chromosome, excluding the pseudoautosomal regions. The median template count is 91 over the autosomes, but half that (44 total) over the X chromosome.

An orthogonal comparison of the relation of template count to copy number can be made at heterozygous loci. In these situations, the sequence context is virtually identical and we expect that the templates from either allele will have the same PCR efficiency and enzymatic representation. Our representation contains 6310 heterozygous SNPs with sufficient coverage to determine the counts for each allele. Since our sample is a normal diploid genome, theory suggests that the template count of one allele should reflect a binomial distribution, B(*N*, *p*), where *N* is the total number of templates and *p* = 0.5. Indeed, the observed allele counts match the expected distribution and show no excess dispersion or deviation from the null expectation. In Supplementary Figure 2B, the histogram depicts the distribution of template counts for one allele at each locus and the black curve shows the theoretical expectation assuming the template counts match the observed.

### Sequence fidelity

As with other methods in which initial templates are tagged, muSeq can be used to reduce sequence error (10,11). Multiple independent reads derived from the same template should consistently support the same sequence; inconsistency is evidence of error. To provide a measure of fidelity, we restricted attention to positions in the genome that are known to be homozygous in the donor and in agreement with the reference genome. Looking at well-covered templates, with 20–100 reads and in which the clustering by conversion pattern was confident, we recorded the proportion of total reads at each position that match the reference base (“reference-base-ratio”). Restricting to such positions, this yields information from 200 million template positions covered by a total of 6.4 billion read bases. We plotted the frequency distribution of the reference base ratio for bit (red) and non-bit (blue) positions (Figure 3). As expected, the reference-base ratios at known homozygous bit positions are bimodal, with 60% of bases unconverted (ratio near 1) and 40% (ratio near 0). For non-bit positions, we set a consensus rule that a cluster ‘reports’ a base position when at least 80% of reads are in agreement. With that rule, 99.93% of high confidence clusters reported the reference base at the previously determined homozygous reference positions. The method, therefore, allows unambiguous recovery of both bit and non-bit genomic sequence without introducing significant new variation.

**Figure 3:**
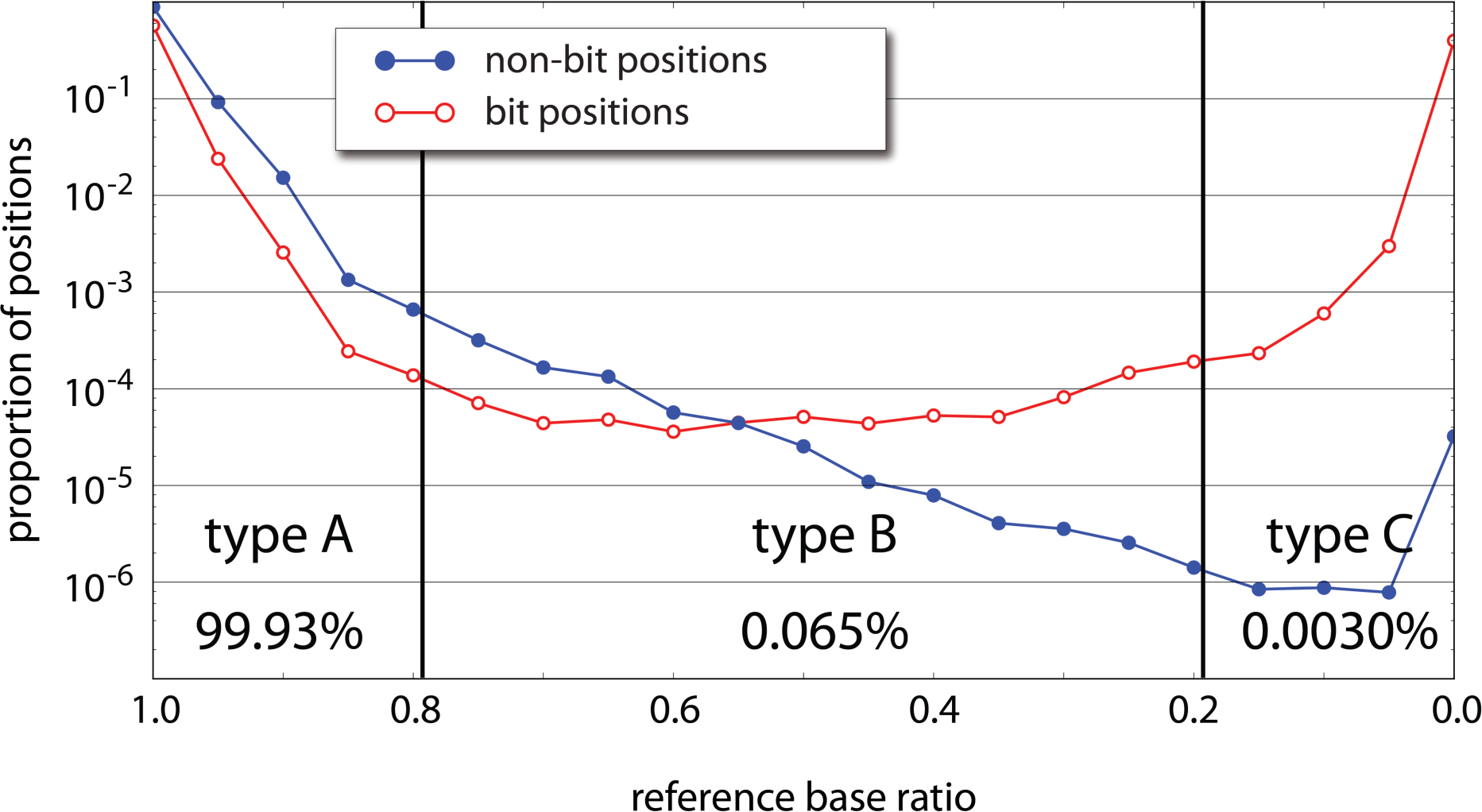
Sequencing error rates. At genomic positions where our sample is homozygous for the reference base, we expect all reads to report the reference base unless (1) the position is a C that converts to T, (2) there is machine error in the read, (3) there is an error introduced during PCR, (4) the initial template is damaged, or (5) the template records a rare somatic variant. From 200 million homozygous template positions, we record the proportion of reads from template clusters (with at least 20 reads per cluster) reporting the reference base (“reference base ratio”, x-axis). The y-axis is a log-scaled normalized histogram of reference base ratios for both bit positions (red) and non-bit positions (blue). Bit positions show a bimodal distribution with a 60:40 split of unconverted to converted positions. Among non-bit positions, 99.93% confidently report the reference base with a ratio greater than 0.8 (type A). Another 0.065% of positions have an uncertain consensus within the cluster with the reference base accounting for between 20% and 80% of reads (type B). About 0.0030% of reads are confidently non-reference (type C). Patterns of base substitutions between these three types (Supplementary Figure 3) suggest that type A are primarily machine error, type B are primarily PCR error, and type C are somatic mutation and/or initial template damage.

Among the non-bit positions, 99.93% had a reference base ratio greater than 0.8. We refer to these as “type A” positions. Positions with a reference base ratio between 0.2 and 0.8 are called “type B” and accounted for 0.065% of non-bit positions. Lastly, 0.003% of non-bit positions had at least 80% reporting a non-reference base. These comprise “type C” positions. Type A and type B positions occurred at rates consistent with machine error and polymerase error, respectively, and each is consistent with a template-error symmetry signal consistent with that interpretation (Supplementary Figure 3). The incidence of type B falls off in log-linear fashion, until there is a rise in incidence of type C. Type C positions could be the result of somatic variation, first-round synthesis polymerase error, or template damage. These have different expected symmetry signals, but from the actual observed symmetries, we infer that type C positions arise mostly from template damage.

### Transcript assembly from cDNA

We next tested whether muSeq could aid in transcript assembly when applied to cDNA. cDNAs were prepared from a fibroblasts using conventional oligo(dT) and template-switching primers that were modified to include random barcodes and bisulfite-resistant PCR primers. After reverse transcription, cDNA was split into three equal batches. The first batch was prepared for libraries in the usual manner: PCR amplified, sheared, and adapted with batch-specific (barcoded) Illumina sequencing primers. The second and third batches were first treated with the muSeq protocol for 40% conversion, and then libraries were prepared with different batch-specific primers. To survive the bisulfite conversion reaction, the custom primers lacked G, and additionally all cytosines in the 3′ primers were methylated. The batches were then pooled together, sequenced, and mapped to the genome. The two muSeq libraries and the unconverted library had similar coverage.

For this application, we used a simplified clustering algorithm (Materials and Methods). Reads were mapped using a reduced alphabet, then partitioned by chromosome and strand into non-overlapping connected components. Each component was clustered using only the expected bit positions within annotated transcripts, followed by applying a greedy algorithm. Although not as exhaustive as the previous clustering algorithm, the clustering was robust. Clusters rarely contained mixed reads from separate batches (Supplementary Figure 4), generally had no more than a single barcode at the 5′ and 3′ ends, had consistent nucleotides at heterozygous positions, and heterozygous sites separated by at least 150 bp were in the proper phase (Supplementary Figure 5).

A study of the alignment file annotated with cluster index and tag information (available for download at <u>http://wigserv2.cshl.edu/web/museq/)</u> shows many genes are well-covered and properly clustered with a consistent conversion pattern spanning multiple read lengths. To provide some measure of success in assembly, we examined the maximal length per gene for all ‘proper and complete’ clusters. These are clusters that 1) have conversion patterns that match the complementary strand of the gene transcript, 2) contain unique barcodes for both the 5′ and 3′ ends, and 3) have properly oriented 5′ and 3′ ends.

Each gene is represented by a single point in the scatter plot (Figure 4A). Average lengths of the annotated transcripts are on the x-axis, and maximal length in muSeq clusters is on the y-axis. For those genes with non-trivial coverage in muSeq and average annotated transcript lengths between 300–6000 bp, 47% have muSeq clusters greater than the average transcript length and 79% are within half the average transcript length. This comparison is intrinsically noisy, as the longest transcripts present in our sample may not reflect the average transcript. As an illustration, we highlight three specific genes RPS15A, RP11-1035H13.3 and ARL6IP1 (Figure 5, green, red and blue, respectively). The RPS15A gene has many annotated transcripts with unspliced introns that are longer than the observed muSeq transcripts. The RP11-1035H13.3 has exactly one known transcript and our one observed cluster covers every annotated position. Lastly, the longest ARL6IP1 muSeq cluster matches the longest known transcript of that gene. Figure 5B displays the distribution of lengths for genes in both muSeq and GENCODE. The distributions for the longest observed muSeq cluster (green) and the average annotated transcript length (blue) are shown. The distributions are well-matched, with a slight skew and a longer tail to the annotated transcripts.

**Figure 4:**
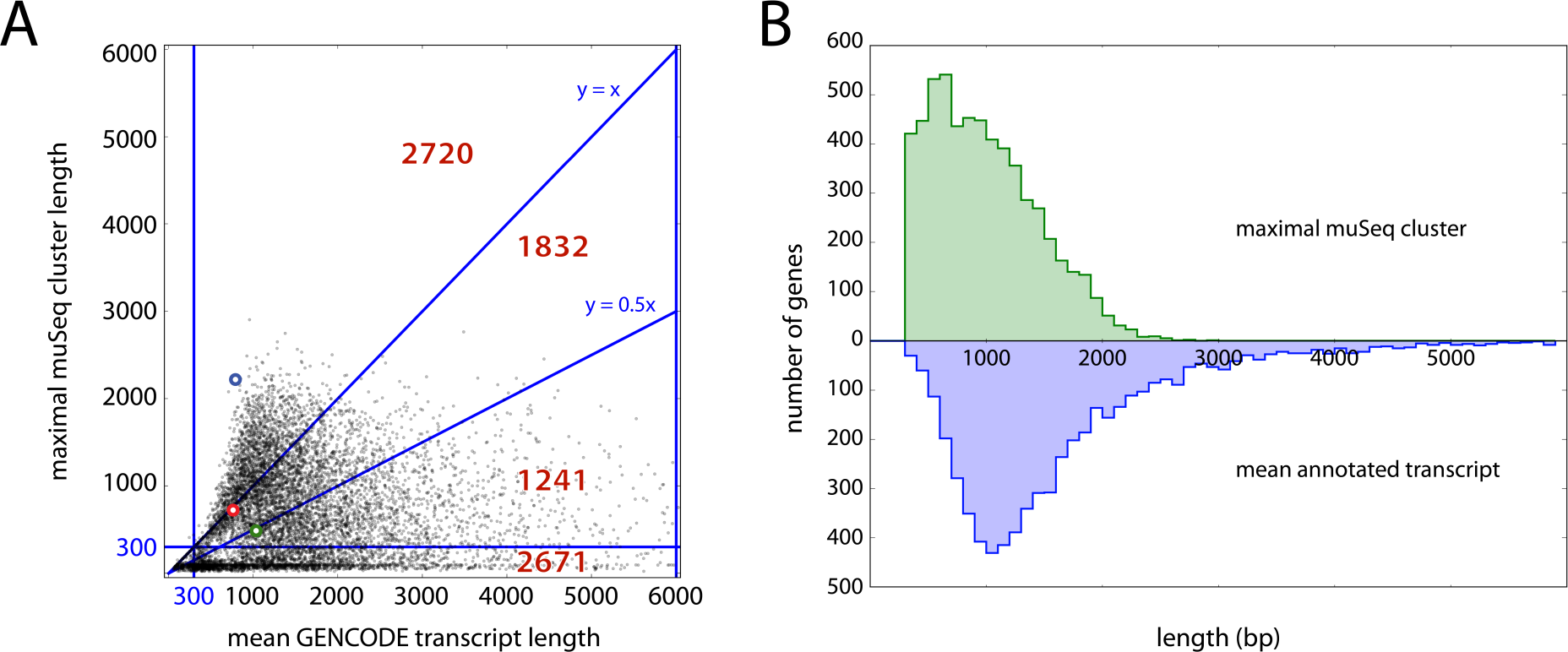
muSeq cluster length compared to annotated genes. **(A)** For each gene, we compared its average annotated isoform length (x-axis) with the longest ‘proper and complete’ muSeq cluster covering it (y-axis). The plot is subdivided into regions with red numbers indicate the number of genes in each region. The highlighted blue, green and red circles correspond to the genes ARL6IP1, RPS15A and RP11-1035H13.3, respectively (described in detail in Figure 5). The longest transcript for ARL6IP1 is observed in the muSeq library, and its length far exceeds the average transcript length (blue dot above y = x). We also observe the full length of the only annotated RP11- 1035H13.3 transcript in the muSeq library (red dot on line y = x). Whereas we observe many full-length muSeq transcripts of RPS15A, the GENCODE database includes many long, unspliced isoforms that are not present in the muSeq library (green dot on line y = 0.5x). **(B)** A histogram of the maximal muSeq cluster length per gene is shown in green, and the average length over all GENCODE-annotated transcripts of that gene in blue. The distributions are well-matched, with a slight shift and heavier tail to the annotated transcripts.

Next, to determine if clusters detect distinct splicing patterns, we surveyed genes with at least one proper and complete cluster >500 bp in length. There are approximately 5000 such genes (detailed plots for each are included in Supplementary Data). The plots elide over intergenic and intronic regions and indicate direction of transcription. Some genes express a range of isoforms, with different exon skipped, varying transcription start and termination sites, and transcripts that do not conform to known annotations. We use one example (Figure 5) to explain the plots and to illustrate the richness of the observed transcript variation. The transcribed regions from two adjacent genes are displayed, with gray vertical stripes denoting compression of intergenic and intronic regions. *ARL6IP1* (blue) and *RPS15A* (green) have ranges of known isoforms depicted in the lower half of the figure and one hybrid transcript (RP11-1035H13.3, red) that includes exons from both genes. The upper panel shows all proper and complete clusters in the muSeq data. Clusters are colored to best match the annotated gene. The *ARL6IP1* clusters are very similar to each other and well-matched to known transcripts, but with a small offset in the 5′ start position and some variability at the 3′ end. *ARL6IP1* is well-covered with a muSeq cluster of 2253 bp and a longest annotated transcript of 2409 (Figure 4A). The observed transcripts of *RPS15A* differ more from the annotated transcripts, having a few variations in the 5′ start position, a variant length of the first exon, and two major variants at the 3′ end. *RPS15A* has a longest muSeq cluster of 529 bp, and the longest annotated transcript is 3828 (Figure 4A). However, the latter is due to an unspliced form that was not observed in this cell line.

**Figure 5.**
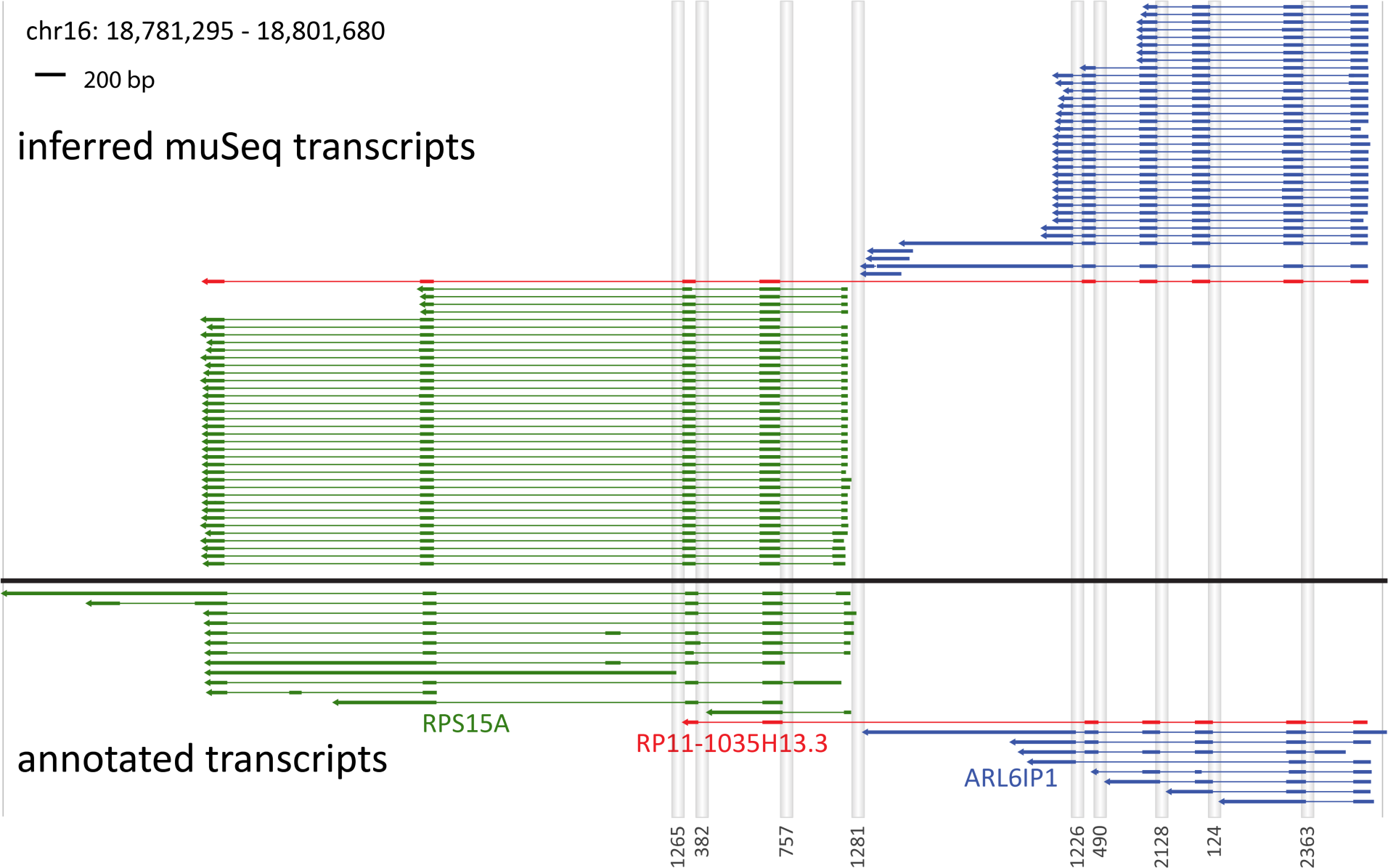
Coverage from proper and complete clusters. Annotated and inferred muSeq transcripts for genes in a 20-kb region of chromosome 16 are shown. There are three annotated genes in the region: ARL6IP1 (blue) and RPS15A (green), as well as the fusion gene RP11-1035H13.3 (red). Genomic segments that are intronic or intergenic are compressed as grey columns labeled by length at the bottom. The lower plot (below black line) shows transcripts of the genes that appear in the ENCODE database. Thick lines are exons and UTRs, thin lines are introns, and the arrows point 5′ to 3′. The upper plot (above the black line) shows the inferred transcripts from the observed muSeq clusters. The clusters shown have consistent tags at both ends of the assembly and conversion patterns that match the strand of transcription. Each cluster is colored to match the most similar annotated gene. Both ARL6IP1 and RPS15A have one major splice pattern with variability in the 3′ and 5′ ends. RPS15A also shows a minor splice variant for the first to second exon junction (lowest four clusters). Additionally, there is a novel transcript that matches the fusion gene RP11-1035H13.3, but includes an additional two exons from RPS15A.

Finally, we observe a cluster (red) that spans exons from both genes much like RP11-1035H13.3, and that is predicted to encode a fusion protein that skips the last coding exon of *ARL6IP1*. Unlike the known annotation, however, this cluster extends an additional two exons to the common end of *RPS15A*. While identification of the hybrid splice junction would be possible from unassembled reads, the full length of the transcripts and the deviation from the GENCODE annotation could not be properly inferred from a standard, short-read protocol.

## DISCUSSION

Previously, using simulation and theory, we demonstrated that prior mutation of initial templates could enable new sequencing applications, such as accurate count and assembly. Here we realize the promise of that theory. We show that by minor modification of existing protocols for sodium bisulfite conversion, we can generate partial conversion patterns with a tunable rate. Mutations are uniform and randomly distributed. After amplification and fragmentation, the conversion patterns emergent from sequencing and mapping unambiguously identify the initial template. This property enables a range of powerful applications including accurate counting, low-error sequencing and long-range assembly, all from relatively inexpensive, short-read sequencing.

Our experiments on genomic representations demonstrate that the mutation rates are tunable within the range of 20–60% conversion for unmethylated cytosines. Moreover, the conversions observed are consistent with an independent probability of mutation at each convertible position. We did observe a slight overabundance of completely unconverted templates beyond the expectations of methylation and chance. This may be due to factors such as insufficient mixing, protein-DNA complexes, or contamination from unconverted template. Additionally, we examined the mutational profile of all positions and found no evidence of increased template damage or PCR error as a result of the conversion process, suggesting that the resulting sequence is highly consistent with the true template at all but the convertible positions.

In our theoretical work and simulations, we employed simple algorithms for template identity and assembly. These algorithms assumed that the reads had no errors and were perfectly mapped to a reference genome. Analyzing experimental data required new methods to cluster and assemble actual sequencing data. To cluster reads from the same template that differ by sequencer and PCR error, we developed ‘transitive propagation,’ a novel clustering algorithm designed to handle multi-scale differences: reads in the same cluster differ according to an error rate (≤1%), whereas reads in different clusters differ according to the conversion rate (50%). We direct the interested reader to the transitive propagation manuscript (6).

To demonstrate sequence assembly, we chose the important application of transcriptome profiling. We designed a preliminary computational pipeline applied to cDNA that leveraged methods developed in the genomic representation. These methods first map reads to the reference genome and identify convertible positions, then use the match/mismatch of reads across converted positions to score the likelihood that pairs of reads derive from the same template. The few failures in the cDNA assembly were most often attributable to mis-mapping of oligo(dT) primers to CT dinucleotide repeats in the genome. Our procedure is capable of generating assemblies up to 3 kb in length from reads that are 150 bp long. An atlas of well-covered transcript assemblies, many showing non-canonical splicing patterns, are included in the Supplement.

We are presently developing tools that can assemble initial templates even in the absence of a reference genome or reference transcript map. This will be useful not only for *de novo* assembly of genomic loci and whole genomes, but also for making precise transcriptome maps from known organisms absent prior knowledge.

## AVAILABILITY

1) Mapped and clustered sequence files for cDNA data and 2) a compilation of 4975 genes with one ‘proper and complete’ cluster >500 bp in length (including depictions of the muSeq transcripts observed in the neighbourhood of each gene) are available for download at http://wigserv2.cshl.edu/web/museq. The directory includes a “read_me” file with details on the user-specified .bam fields. The format of each plot in the compilation mirrors that of Figure 5. Known transcripts are in color and labeled, whereas the muSeq sequences are in black.

## SUPPLEMENTARY DATA

Supplementary Figures 1–5 are available as a single .docx file **at NAR online**.

## ACKNOWLEDGEMENT

We thank Peter Andrews, Alex Dobin, Jude Kendall, Bud Mishra and Michael Schatz for assistance, suggestions and feedback throughout. The authors wish it to be known that, in their opinion, the first 2 authors should be regarded as joint First Authors.

